# Decoding interaction-induced proteome changes in co-cultures with hybrid quantification and SILAC-directed real-time search

**DOI:** 10.1101/2025.09.09.675040

**Authors:** Alexia Carré, Sofía Ibáñez-Molero, Daniel S. Peeper, Maarten Altelaar, Kelly Stecker

## Abstract

Intercellular communication between T cells and cancer cells plays a pivotal role in determining cancer cell survival or death. Yet, our understanding of this interaction remains incomplete. Methods to study heterotypic cell interactions are either limited to targeted studies relying on predefined set of proteins, or require cell separation, thus disrupting the native environment.

Stable isotope labeling by amino acids in cell culture (SILAC) enables proteome distinction in heterologous co-cultures without the need for physical separation. But dynamic studies remain constrained by the need for numerous mass spectrometry (MS) runs, the challenges in detecting low-abundant proteins, particularly in immune cells and the limited data completeness due to the use of data-dependent MS^1^-based precursor quantification.

To overcome these limitations, we evaluate the integration of SILAC with tandem mass tag (TMT) multiplexing and SILAC-directed real-time search (RTS). TMT labeling enables simultaneous analysis of multiple samples, while RTS-MS^3^ acquisition using SILAC-induced mass shifts as fixed modifications triggers MS^3^ scans for specific proteome populations within a mixed cell system, improving quantitative accuracy and proteome coverage for target protein populations.

We benchmarked our acquisition methods using SILAC-labeled samples mixed at defined ratios and validated the approach in biologically relevant co-culture experiments. Additionally, we introduced a carrier channel to enhance detection of lower-abundant T cell proteins, while maintaining acceptable quantitative precision. Our results demonstrate that the combined SILAC-TMT-RTS strategy dramatically improves proteome depth, temporal resolution, and cell-type specificity for short-term co-culture interaction proteomics studies. In co-culture samples of T cells with non-small cell lung cancer cell lines that were either sensitive or resistant to T cell killing, our method revealed candidate mechanisms underlying their differential sensitivity. Our integrated approach combining SILAC, TMT and RTS to resolve cell-specific proteome dynamics in co-culture represents a novel and powerful advance.

## Introduction

Understanding protein dynamics in interacting immune and cancer cell populations is crucial for elucidating intercellular communication and underlying disease mechanisms on a molecular level. However, methods to comprehensively study these interactions at the protein level are limited. Techniques such as mass cytometry or immunohistochemistry typically rely on the analysis of a predefined set of proteins thus hindering extensive and discovery-based characterization. In contrast, standard proteomics offer broad characterization but require physical separation of interacting cells via fluorescence-activated cell sorting (FACS), which disrupts the cellular context and can be especially challenging for cells engaged in strong physical contact, such as T cells and tumor cells that form immune synapses.

Stable isotope labeling by amino acids in cell culture (SILAC) has traditionally been used to compare experimental conditions by mixing labeled proteomes before or after cell lysis (1). SILAC has also been adapted for dynamic studies, including pulsed-SILAC to investigate protein turnover, (2–4), and more recently for probing both distant and proximal cellular communication (5, 6). Combined with proximity labeling, SILAC has enabled the surface proteome profiling of cell-cell interface in co-culture (7). SILAC also provides a solution for the analysis of mixed proteomes from interacting cells. By labeling each cell population with distinct isotopically labeled amino acids, prior to co-culture, cells can be collected at once and proteomes untangled after mass spectrometry (MS) acquisition (8, 9). We recently applied such a strategy to the co-culture of tumor and T cells, and termed this strategy HySic (for hybrid quantification of SILAC-labeled interacting cells) (10).

Despite its advantages, SILAC-based dynamic proteomics face practical limitations. First, each timepoint requires a separate MS run, making large-scale temporal studies costly and time-consuming. Second, data-dependent acquisition (DDA) studies using MS^1^-based quantification approaches suffer from missing data, making the interpretation of protein dynamics difficult (11). The challenge is further exacerbated in co-culture systems with immune cells. Given that T cells are smaller than tumor cells (5-7 µm vs > 10 µm) (12, 13), they contain less protein, making their detection challenging especially considering the stochastic nature of DDA, which tends to undersample low-abundant proteins.

Tandem mass tags (TMT) offer a promising solution to these challenges. These isobaric reagents are widely used for sample multiplexing and their reporter ion intensities are reflective of relative protein abundance across samples. TMT has been successfully combined with SILAC in hyperplexing strategies and for studying proteoform and protein dynamics (14–16). Significant technological improvements have established that using a dedicated quantitative spectrum for reporter ion quantification (MS^3^) together with synchronous precursor selection (SPS) offer the highest accuracy (17, 18). However, the additional third scan required for MS^3^ slows the acquisition speeds, thereby leading to lower proteome coverage. The integration of a real-time search (RTS) which triggers the acquisition of quantitative MS^3^ scans only upon confident peptide identification has demonstrated increased acquisition speed restoring MS^2^-level proteome coverage while maintaining accurate quantitative measurements (19, 20).

For low abundant samples such as single-cell proteomics, the addition of a carrier proteome in one of the TMT channels has been successfully and consistently used to boost the sensitivity and to enable the identification and quantification of more peptides (21–23). But this approach suffers from a loss in quantitative accuracy resulting from overloading the analyzer with the carrier channel ions.

In this study, we evaluate the combination of TMT multiplexing with SILAC-labeled interacting cells. We first provide a detailed characterization of proteome coverage and quantitative accuracy in SILAC-labeled samples mixed at different artificial or biological-mimicking ratios using MS^2^, SPS-MS^3^ or RTS-MS^3^ acquisition methods. We also assess the feasibility to use SILAC-induced mass shifts as a fixed modification for RTS to trigger MS^3^ scans. We further demonstrate the benefit of this improved strategy in co-culture experiments by characterizing proteome changes in T cells, tumor cells and newly-translated proteins across two non-small cell lung cancer (NSCLC) cell lines, one sensitive (NCI-H358) and one resistant to T-cell killing (A549). Our findings suggest that combining TMT with SILAC and RTS enhances proteome depth and temporal resolution while preserving sufficient quantitative accuracy to support novel biological insight.

## Experimental procedures

### Experimental design and Statistical rationale

A description of each relevant experimental design is available in Fig 1A., 2A and S4A. The data acquired for this study was generated from replicates mixed samples multiplexed with TMT or TMTpro reagents and one to two MS acquisitions for method evaluation (artificial mix of two distinct SILAC-labeled proteomes). Controls consisted of multiplexed single proteome for comparison of quantification accuracy and protein/peptide counts. For co-culture samples, two to three biological replicates were multiplexed with TMTpro reagents and one MS acquisition was carried for each RTS setting. Datasets were analyzed with MSstat TMT for statistical evaluation. Significantly changing proteins are included in the analyzes if adjusted p-value < 0.05. All raw peptides or PSMs output from PD can be found in Supplementary Tables.

**Figure 1.**
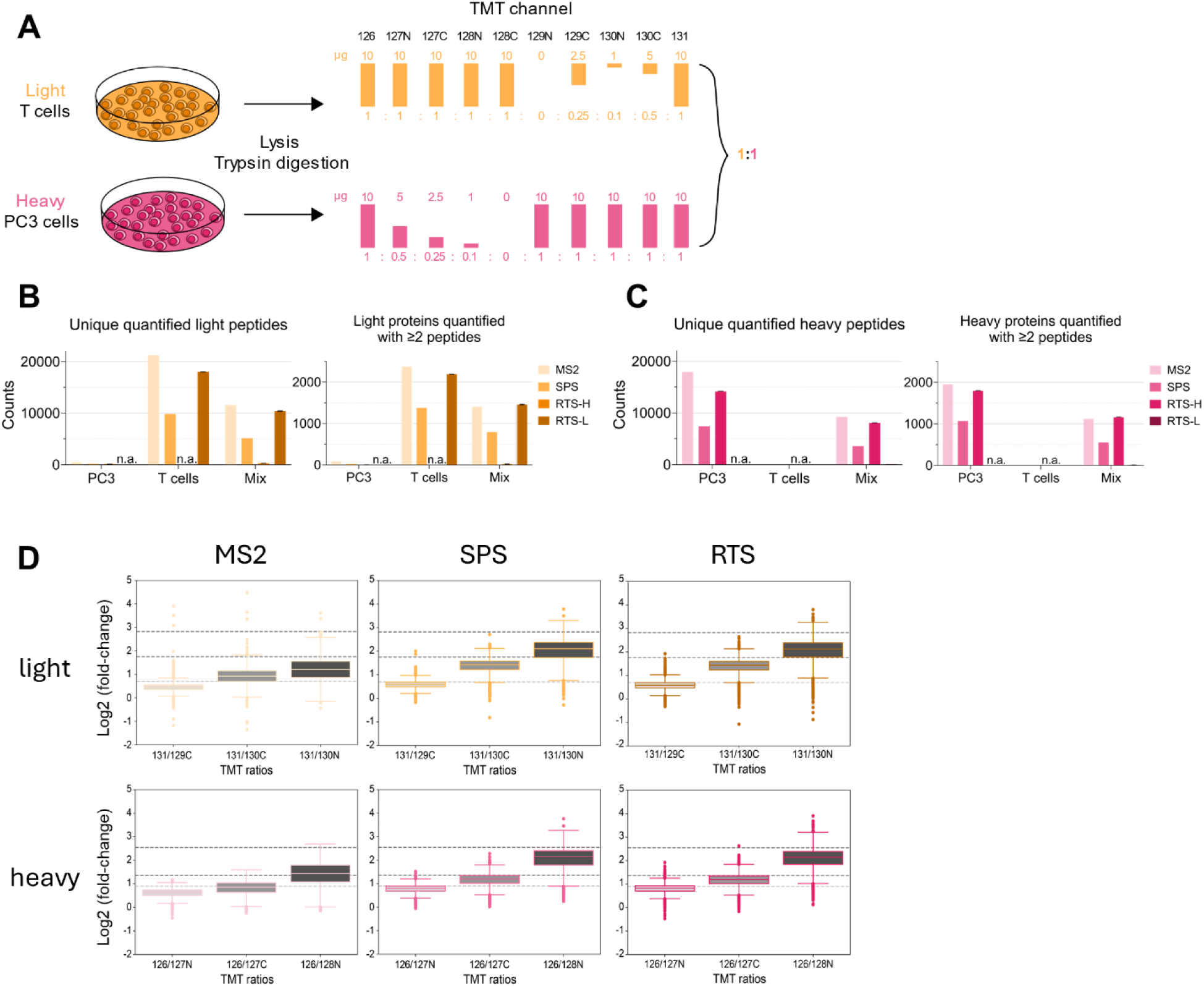
Evaluation of TMT implementation in mixed SILAC samples at 1:1 ratio. **(A)** Light-labeled T cells and heavy-labeled PC3 tumor cells were lysed, digested and TMT-labeled prior to being mixed so that the total of heavy and light content is equal across TMT channels while still allowing various ratios within the same SILAC label. **(B)** Light and **(C)** heavy peptide and protein counts across samples (PC3 cells alone, T cells alone or mixed sample) and acquisition strategies. n.a.: not applicable, n=1 injection for SPS and MS2, n=2 injections for RTS acquisitions **(D)** Distribution of quantified light (top) and heavy (bottom) peptides using TMT ratios across relevant channels and across MS acquisitions. Best measured ratio as observed in Fig. S1 are represented by horizontal dashed lines.

### Cell culture and SILAC labeling

NK-92 cells were maintained in RPMI (Capricorn Scientific, RPMI-STA) supplemented with 10% FBS (Gibco, A5256701), 100U/mL Penicillin/Streptomycin (Gibco, 15140-122) and 100U/mL recombinant human IL-2 (Peprotech, HZ-1015). Cells were collected by centrifugation (100g, 5 min), washed twice with PBS (Capricorn Scientific, PBS-1A) and dried pellets were frozen at −80°C until further use. PC3 cells were cultured in SILAC RPMI (Gibco, 88365) supplemented with 10% dialyzed FBS (Gibco, 89986), Penicillin/Streptomycin, 2 mM Glutamax (Gibco, 35050061), 0.1 mg/ml L-Arginine:HCL (13C6;15N4) (Cambridge Isotope Laboratories, CNLM-539-H-0.25), 0.04 mg/ml L-Lysine:2HCL (3,3,4,4,5,5,6,6-D8) (Cambridge Isotope Laboratories, DLM-2641-PK), 0.2 mg/ml L-proline (Sigma-Aldrich, P5607). Cells were detached with trypsin-EDTA (Gibco, 25300054), washed with PBS and frozen as dry pellets.

### Sample lysis and digestion

Cell pellets of 0.2-1×10^6^ NK-92 or PC3 cells were resuspended in 50µL of 5% sodium dodecyl sulfate (SDS) in 50 mM triethylammonium bicarbonate pH 8.5 containing a protease inhibitor (Roche, cOmplete Mini EDTA-free, 11836170001) and sonicated with 10 cycles (30s on, 30s off) in a Bioruptor Plus (Diagenode). Lysates were clarified by centrifugation at 13,000g for 10 min at 4°C. Protein concentration was measured using a BCA protein assay (ThermoFisher Scientific, 23225). Samples were digested for 2h at 47°C with trypsin (1:15 enzyme to protein ratio, Sigma-Aldrich) using S-trap mini (Protifi, C02-mini-80) or S-trap micro columns (Protifi, C02-micro-80), following manufacturer’s instructions. Eluted digested peptides were dried down in a vacuum centrifuge prior to TMT labeling.

### HySic samples

Co-culture and T-cell digested and desalted samples were obtained from a previously published study (10). Briefly, A549 or NCI-H358 tumor cells were transduced with the HLA-A∗02:01-MART1-mPlum lentiviral plasmid and SILAC-labeled with heavy arginine (^10^R) and lysine (^8^K). Primary T cells from 3 different donors were transduced with a MART-1-specific TCR. Tumor and T cells were co-cultured in medium containing “intermediate” arginine (^6^R) and lysine (^4^K) for 0h, 4h or 6h.

### TMT and TMTpro labeling

10-15µg of digested sample were resuspended in 20µL of 20% (v/v) acetonitrile (ACN) in 50mM HEPES buffer and labeled with 4µL of TMT 10-plex reagents (ThermoFisher Scientific, 90110) at a label to protein ratio of 4:1 for 1.5h at 300rpm, at room temperature. Alternatively, 10-15µg of digested sample were resuspended in 20µL of 50mM HEPES buffer and labeled with 4µL of TMTpro 18-plex reagents (ThermoFisher Scientific, A52047 and A52048) at a label to protein ratio of 5:1, for 1h at room temperature. T cell digests of the 3 donors were mixed in a 1:1:1 ratio prior to labeling, to minimize donor-dependent observations. Labeling efficiency and mixing ratios were checked by mixing 1uL of each TMT(pro) labeling reaction in an acidic solution and running 250-500ng in a liquid chromatography tandem mass spectrometry (LC-MS/MS) run. Once labeling was confirmed to be >95%, reactions were quenched with 2µL of 5% hydroxylamine in 50mM HEPES and incubated for 15min at room temperature. Labeled samples were mixed according to mixing schemes (Fig. 1A, 2A and S4), dried down in a vacuum centrifuge and desalted with either Oasis HLB plate (Waters, 186001828BA) or SepPak C18 cartridges (Waters, WAT054960). Samples were vacuum-dried and kept at −80°C until use.

The HySic mixed samples (Fig. S4) underwent high-pH fractionation. Each sample was resuspended in 20µL of 10mM of NH_4_OH pH 10 (Buffer I) and eluted at a flow rate of 200µL/min on a Kinetex 5µm EVO C18 150 x 2.1mm column (Phenomenex, 00F-4633-AN) with a gradient of 90% ACN in 10mM NH_4_OH pH 10 (Buffer II). The gradient was as follow: 0-2% buffer II for 2 min, 2-12% for 6 min, 12-35% for 47 min, 35-55% for 7 min followed by a wash and equilibration phase. Fractions of 1 min were collected from minute 1-75. Non-adjacent fractions were combined in 8 different pools. The pools were dried in a vacuum-centrifuge and kept at −80°C until use.

### LC-MS/MS acquisition

TMT(pro)-labeled samples were resuspended in 2% formic acid (FA) in water and analyzed on an Orbitrap Eclipse Tribrid Mass Spectrometer (ThermoFisher Scientific) coupled to an Ultimate 3000 UHPLC (ThermoFisher Scientific). Peptides were trapped on a pre-column (PepMap Neo 5 µm C18 300 µm x 5 mm Trap Cartridge, ThermoFisher Scientific, 174500) for 1 min in 4% buffer B (0.1% FA in 80% ACN), 96% buffer A (0.1% FA). Peptides were then separated on an in-house packed analytical column (ReproSil-Pur 120 C18-AQ, 2.4 µm 75 µm x 50 cm for the 1:1 mix and ReproSil-Pur 120 C18-AQ, 1.9 µm 75 µm x 50 cm for the other samples, bulk media from Dr. Maisch) with a gradient from 4% to 13% buffer B in 2 min, from 13% to 40% of B in 94 min, from 40% to 55% in 15 min, from 55% to 99% in 1 min followed by 5 min of wash at 99% of buffer B and equilibration (from 99% to 4%) in 15 min. Flow rate was set to 300 nL/min.

All MS acquisitions were data-dependent acquisitions with slight variations depending on the method. Unless otherwise stated, MS^1^ scans were acquired in the Orbitrap with a resolution of 120,000, scan range of 400-1600, normalized AGC target of 100%, injection time mode set to auto and intensity threshold to 5×10^3^. Precursors with charge state 2-6 were selected for fragmentation with an isolation window of 1.2 Th and a dynamic exclusion of 60s.

For MS^2^-based acquisition, the cycle time was set to 3s and MS^2^ scans were measured in the Orbitrap with a resolution of 50,000, HCD collision energy of 38% (TMT) or 35 (TMTpro), isolation window of 0.7 Th, first mass of 120 m/z, max injection time of 200ms and normalized AGC target of 250% (TMT) or 200% (TMTpro). MS^1^ precursor intensity threshold was 2.5×10^4^.

For SPS acquisitions, the cycle time was set to 3s and MS^2^ scans were acquired in the IonTrap with a scan rate set to turbo using HCD collision energy of 36% (TMT) or HCD collision energy 35% (TMTpro), isolation window of 0.7 Th, scan range of 400-1600, max injection time of 35 ms and normalized AGC target of 100%. MS^3^ scans were acquired in the Orbitrap with a resolution of 50,000, scan range of 100-500m/z, a normalized AGC target of 200%, max injection time of 200ms, MS isolation window of 0.7 Th, MS^2^ isolation window of 2 Th. 10 SPS precursors were fragmented with HCD and a collision energy of 65 (TMT) or 55 (TMTpro).

For RTS acquisitions, MS^1^, MS^2^ and MS^3^ parameters followed the ones of SPS acquisition with the following adjustment: MS^1^ dynamic exclusion was reduced to 45s, cycle time set to 2.5s (TMT) or 3s (TMTpro). MS^2^ used HCD at 36% (TMT) or CID at 30% (TMTpro). The RTS was carried out on the human proteome fasta file (20,422 entries, downloaded 18 April 2024) with trypsin as the enzyme. The search parameters were Xcorr of 1.4, dCn of 0.1, precursor ppm of 10, charge state of 2 and dynamic exclusion of 60s allowing for max 4 peptides per protein. For all RTS acquisitions, variable modification was methionine Oxidation (+15.9949) and static modification Carbamidomethyl (+57.0215) on cysteines. Depending on the SILAC label of the sample, lysines could be either TMT(pro)-labeled or SILAC and TMT(pro)-labeled. This was taken into consideration during RTS by implementing new modifications. For RTS-L acquisition, static modifications also contained TMT6plex (+229.1629) or TMTpro16plex (+304.2071) on lysines and peptide N-termini. For RTS-H acquisition, heavy-labeled arginines (+10.008) was consistently included as a static modification along with TMT6plex (+229.1629) on peptide N-termini and TMTheavy (+237.2131) on lysines for TMT experiment or TMTpro16plex (+304.2071) on peptide N-termini and TMTproHeavy (+312.2573) on lysines for TMTpro experiments. For RTS-I acquisitions, static modifications consisted of TMTpro16plex (+304.2071) on peptide N-termini, intermediate-labeled arginines (+6.02) and TMTproInt (+308.23225) on lysines.

### MS data processing

Raw data files were processed with Proteome Discoverer 3.1 (ThermoFisher Scientific). For each dataset, two to three parallel Sequest HT branches were configured using a UniProt database (20,433 entries, downloaded 05 December 2024) and a contaminant database (from MaxQuant software). Across all branches, the following settings were applied: enzyme was trypsin, max missed cleaveages were of 2, minimum peptide length was 7, precursor mass tolerance was 10ppm, static modifications included carbamidomethyl on cysteines (+57.021 Da) and TMT (+229.1629Da) or TMTpro (+304.207 Da) on peptide N-terminus and dynamic modification include Oxidation of methionine residues (+15.995 Da). MS^2^ datasets were searched with a fragment mass tolerance of 0.05 Da and MS^3^-based datasets with 0.5 Da. Each Sequest HT node reflected labeling schemes and related mofidications: light label included TMT (+229.1629 Da) or TMTpro (+304.207 Da) on lysines (static), heavy label included TMT-Lys-D8 (+237.2131 Da) or TMTpro-Lys-D8 (+312.257 Da) on lysines (dynamic) and heavy arginines (+10.008Da, dynamic), intermediate label included TMTpro-Lys-D4 (+308.23225 Da) on lysines (dynamic) and intermediate arginines (+6.02 Da, dynamic). The node for intermediate label was only used for HySic samples. Each search node was followed by a Percolator node with FDR targets of 0.01 (strict) and 0.05 (relaxed). Reporter ions were integrated with a tolerance of 20ppm for both MS^2^ and MS^3^-based quantifications. Only unique peptides were used for quantification, no normalization nor scaling were applied. The thresholds of co-isolation, minimum average reporter S/N, SPS Mass matches and site probability were respectively 75 (MS^3^) or 50 (MS^2^), 10, 65% and 75. The peptide- and protein-level FDR was 1%.

### MS data analysis

For all analyzes, contaminants were excluded and only unique sequences were considered (assigned to one protein). For performance evaluation in 1:1 and 1:5.5 mixes, peptide exports from Proteome Discoverer were used. Proteins were retained only if at least 2 peptides were identified. For HySic samples, intensities in the PSM output were median normalized using scaling factors derived from SPS acquisitions of the unfractionated sample. PSMs were then filtered (Fig. S3B) and fed to MSStat TMT (v2.14) for protein quantification and statistical analyzes (24).

For heatmap analysis, tumor and T cell proteins were further filtered to include only those exhibiting a decrease in both cell lines (6 h and 4 h ≤ 0 h), with statistical significance (adjusted p-value ≤ 0.05) in at least one cell line and one time point compared to 0 h. Heatmap analysis of newly-translated proteins was performed on proteins with a significant 1.8-fold increase (i.e. a 2-fold corrected for the observed ratio compression) in either cell line (6h or 4h > 0h, adjusted p-value ≤ 0.05). Within each cell line, protein intensities were normalized to the average intensity at 0 h.

For volcano plot visualization, proteins were filtered as described above for upregulated (newly-translated) or downregulated (tumor and T cells) proteins, prior to selecting the comparison of interest (6 h vs 0 h). Fold-change cutoff was set to log2(1.8).

Metascape (25) was used for pathway enrichment analysis and included Reactome, Wikipathways, KEGG and Hallmark. Significantly enriched pathways (p < 0.01) were identified by comparing the proteins of interest to appropriate background sets: all identified proteins carrying the same SILAC label, combined with newly synthesized proteins for tumor and T cell analyses, and all identified proteins (across all SILAC labels) for newly synthesized protein analysis.

All plots were generated with GraphPad Prism (v10) or custom R scripts.

## Results

### Performance assessment in a 1:1 mixed SILAC proteome

To benchmark the performance of different acquisition strategies in a balanced TMT-SILAC context, where mixed cell types contribute equally to the overall protein content, we compared MS^2^-based acquisitions to SPS-MS^3^ (“SPS” hereafter) and SILAC-directed RTS-MS^3^ (“SILAC-RTS” hereafter) using a 1:1 mixture of light and heavy SILAC-labeled peptides.

Light-labeled primary T cell digests and heavy-labeled PC3 tumor cell digests were TMT-labeled, and titrated across TMT channels in a reciprocal manner. Heavy and light TMT-labeled samples were mixed ensuring a 1:1 peptide ratio of light-to-heavy in the final mixture (Fig. 1A). The mixed sample as well as the separate T cell and PC3 cell samples were analyzed with MS^2^, SPS and SILAC-RTS methods. Two distinct SILAC-RTS methods were designed using the SILAC label as distinguishing modification, with one method targeting light peptides (RTS-L) and the other targeting heavy peptides (RTS-H). In the RTS-H method, heavy SILAC labels were set as static modifications on lysine and arginine. TMT was set to static modification of lysine residues and peptide N-termini in both conditions, resulting in custom TMT-heavy modification on lysines in RTS-H.

We first observed a decrease in peptide and protein counts in all acquisition strategies when comparing the individual cell-type to the mixed cell sample (Fig. 1B, C), emphasizing the challenge of maintaining proteome coverage when investigating complex mixed-proteome samples. As expected, SPS acquisitions suffered from the most pronounced loss in depth, with a 37-59% reduction at both peptide and protein levels relative to MS^2^ and RTS. SILAC-RTS yielded slightly fewer peptides than MS^2^ but, importantly, the number of proteins identified with ≥ 2 unique peptides was identical between the two methods in the mixed sample (∼1,500 light, ∼1,200 heavy). This demonstrates that SILAC-RTS achieves equivalent proteome coverage for each cell type compared to the faster MS^2^ acquisition method, indicating that targeting specific populations with SILAC-RTS in complex samples compensates for longer MS3 duty cycles.

Next, we tested the quantitative accuracy of each acquisition method. To account for technical variability such as mixing errors, we established the true peptide ratios across TMT channels in the T-cell and tumor-cell samples separately (Fig. S1 A, B). These empirical ratios were slightly lower than the theoretical expectations of 1:0.5, 1:0.25 and 1:0.1 (Fig. 1A) and were used as baselines for evaluating quantification accuracy in the mixed sample (Fig. 1D, horizontal grey lines). While in the separate samples MS^2^, SPS and SILAC-RTS methods performed equally, in the mixed sample, MS^3^-based acquisitions displayed a drastically higher accuracy than MS^2^, which was expected based on previous studies (18, 26). The quantification of heavy peptides was more accurate than that of light peptides, possibly due to unlabeled light peptides being present in the heavy samples.

In summary, these results show that SILAC-RTS methods using SILAC labels as static modifications offer a robust solution to achieve deep proteome coverage and high accuracy in 1:1 SILAC-mixed cell, TMT-labeled samples.

### Impact of a carrier channel in an unequal 1:5.5 mixed SILAC-TMT proteome

Immune cells are smaller than cancer cells (12, 13) and for an equivalent cell number lead to 5-6 times less protein content (10). In order to better simulate this proteome imbalance in co-culture samples where the same number of each cell types are incubated together, we designed a mixed SILAC-TMT sample with a final light-to-heavy protein ratio of 1:5.5. Light-labeled NK-92 cells were mixed in various ratios with heavy PC3 cells, with or without the addition of a carrier channel (Fig. 2A). A carrier channel consists of including a large amount of reference sample, in our case, of NK-92 cells, in one TMT channel. This strategy boosts the signal of low-abundant peptides of interest at the MS1 level thereby increasing their detection and quantification.

**Figure 2.**
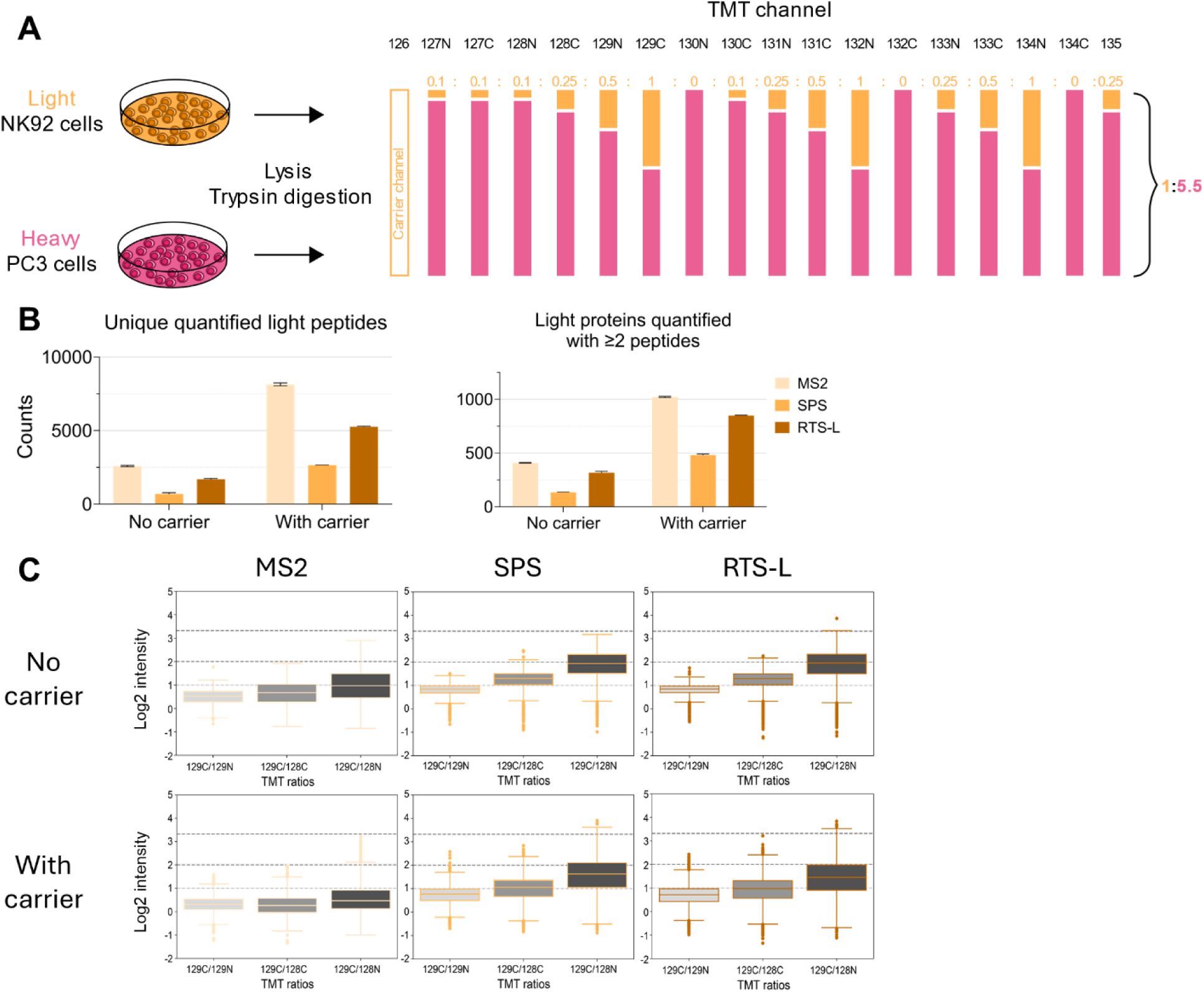
Evaluation of TMT implementation in mixed SILAC samples at a 1:5.5 ratio. **(A)** Light-labeled NK92 cells and heavy-labeled PC3 tumor cells were lysed, digested and TMT-labeled prior to being mixed so that the total of heavy content is 5.5-times the one of light content, while allowing various ratios within the same SILAC label. TMT 126 was left empty or used as a carrier channel (10x the amount of a single TMT channel). **(B)** Quantified llight peptide and protein counts across MS acquisitions in the absence or presence of a carrier channel. **(C)** Distribution of quantified light peptides in the absence (top) and presence (bottom) of a carrier channel. TMT ratios across relevant channels and across MS acquisitions are represented. Expected ratios are shown by the horizontal dashed lines.

In absence of a carrier, the 1:5.5 proteome mix quantified approximately 4-times more heavy proteins than light ones (1,618 vs 410) with MS^2^ acquisition(Fig. 2B, Fig. S2A). This differential detection ratio is expected given the unequal total proteome mixture from the different cells. However, in contrast to equally mixed proteomes, SILAC-RTS acquisition quantified slightly less light-labeled proteins compared to MS2 (Fig. 2B). The low stoichiometry of light peptides limited SILAC-RTS performance relative to MS^2^ due to the reduced frequency of light precursors isolated for MS2 fragmentation, a prerequisite for MS3 RTS-L selection. To address this, we introduced a carrier proteome in the TMT channel 126, containing 10-times the protein amount of each TMT channel, to increase the total amount of light peptide present for MS^1^ precursor selection. This addition significantly boosted protein identifications and achieved a 2.5-fold increase with MS^2^ (1022 vs 410 proteins), 2.7-fold with RTS-L (849 vs 318 proteins) and 3.6-fold with SPS (481 vs 134 proteins), while, concomitantly, heavy protein numbers decreased (Fig. 2A, S2A).

Quantitative accuracy was generally lower in the 1:5.5 mix as compared to the 1:1 mix (Fig. 2C, Fig. 1D). However, SPS and RTS-L acquisitions retained some of the peptide-level expected ratio whereas MS^2^ showed dramatically poor performance with observed ratios substantially deviating from the expected values. Even at the highest mixing ratio, MS^2^ acquisition prevented the detection of variation across channels. To further characterize the effect of the carrier channel, we looked at the distribution of peptide intensities across acquisitions (Fig. S2B). Consistent with previous reports, only the 127C channel was directly affected by the carrier (22), which translated in complete loss of quantitative accuracy in that channel (Fig. S2C). An overall compression of intensity distribution was also noted, which may obscure subtle biological differences.

These findings collectively highlight the importance of designing accurate test samples to reflect the biological context of interest when evaluating a new method. Quantifying TMT ratio compression in samples where the total proteome input is equal for both cell types significantly understimates the problem compared to samples where the proteome content is unequal across cell types. Our method evaluation demonstrates that using light and heavy amino acids independently as fixed modifications in RTS-based acquisitions remains the most effective strategy for the detection and quantification of low-abundant peptides, even in imbalanced mixtures, such as with a 1:5.5 ratio. The use of a carrier channel further enhances peptide identification but involves a trade-off with quantitative accuracy. Therefore, the decision to employ a carrier channel should be guided by the relative importance of identification versus quantitative accuracy in a given sample.

### Development of the data analysis pipeline for low-abundant peptides

Background signal in TMT experiments can make it difficult to accurately quantify peptides, especially low-abundant ones. This challenge is specifically relevant for light-labeled peptides in our co-culture system, which are expected to be present at lower levels due to the smaller size of immune cells.

To better understand the signal characteristics, we examined the light PSMs signal in the 1:5.5 SILAC-TMT mix across acquisition strategies. In all cases, empty TMT channels exhibit an average signal-to-noise (S/N) of 3.2-3.7 (Fig. S3A, left panel). Without a carrier, MS^3^-based acquisitions showed that channels containing only heavy-labeled peptides (e.g., TMT130N, 132C, 134C) had light PSM S/N between 17.2 (RTS-L) and 24.6 (SPS), while channels with minimal light content (e.g., TMT127N, 127C, 128N, 130C) ranged from 23.6 (RTS-L) to 31.1 (SPS). These values suggest that even low-level light signals are distinguishable from background in the absence of a carrier, ensuring an intensity difference of about 6.

However, the addition of a carrier increased background noise, with channels lacking light peptides showing S/N between 7.3 (RTS-L) and 11.0 (SPS), and those with minimal light content averaging 8.5 (RTS-L) to 11.8 (SPS) (Fig. S3A, right panel). This intensity difference of less than 1 makes it difficult to reliably distinguish true signal from background noise. Based on these observations, we established a conservative analysis pipeline. Any measured abundance ≤ 5 was considered background and only PSMs with a S/N ≥ 7 were retained for downstream analysis (Fig. S3B).

In addition to intensity-based filtering, we also closely evaluated variation between observed and expected fold change for the less sensitive condition (2:1 expected ratio). In the 1:5.5 mix, where a log2 fold-change of 1 (i.e., a ratio of 2:1) was expected, we observed a log2 fold-change of ∼0.77 (1.7:1) in the presence of a carrier channel and of ∼0.82 (∼1.8:1) in its absence. To account for this compression and ensure robust detection of biologically meaningful changes, we adopted a fold-change threshold of 1.8 for identifying significantly up- or downregulated proteins.

Altogether, the filtering strategy was designed to enhance quantification reliability particularly for low-abundant peptides. By excluding ambiguous signals and focusing on high-confidence PSMs, we aimed to reduce false positives and improve the interpretability of our proteomics data.

### Combination of TMT with SILAC leads to better coverage and resolution in co-culture samples

To highlight the significance of our approach for biological discoveries, we applied it to co-culture samples from our HySic study (Fig. S4A) (10). In this work, heavy-labeled NSCLC tumor cell lines were co-cultured with light-labeled primary T cells in a medium containing intermediate arginines and lysines, so that all newly-translated proteins carry intermediate labeling. Samples were collected at 0, 2, 4, and 6 hours after co-culture. Out of the initial 4 NSCLC cell lines, we focused on the most resistant and the most sensitive to T cell killing, A549 and NCI-H358, respectively (27). The 2h timepoint was excluded from the present study, as we here focus on proteome-level changes. Replicates were labeled with TMT, pooled and fractionated using high-pH reverse-phase chromatography. All 8 fractions were acquired with RTS methods using either light (RTS-L), intermediate (RTS-I) or heavy (RTS-H) amino acids as fixed modifications. For RTS-L, a 10x carrier channel of primary T cells digests was used and TMT 127C was left empty.

After normality assessment of each dataset, the data were analyzed with MSstats TMT (24) (Fig. S3B, S4B-D). RTS-H quantified 70,583 unique PSMs and 38,561 peptides, RTS-I 16,435 PSMs and 9,933 peptides and RTS-L 97,976 PSMs and 27,416 peptides (Fig. 3A). Each RTS method showed high specificity, with low quantification rate of the other labels (e.g. few intermediate or light PSMs/peptides in RTS-H).

**Figure 3.**
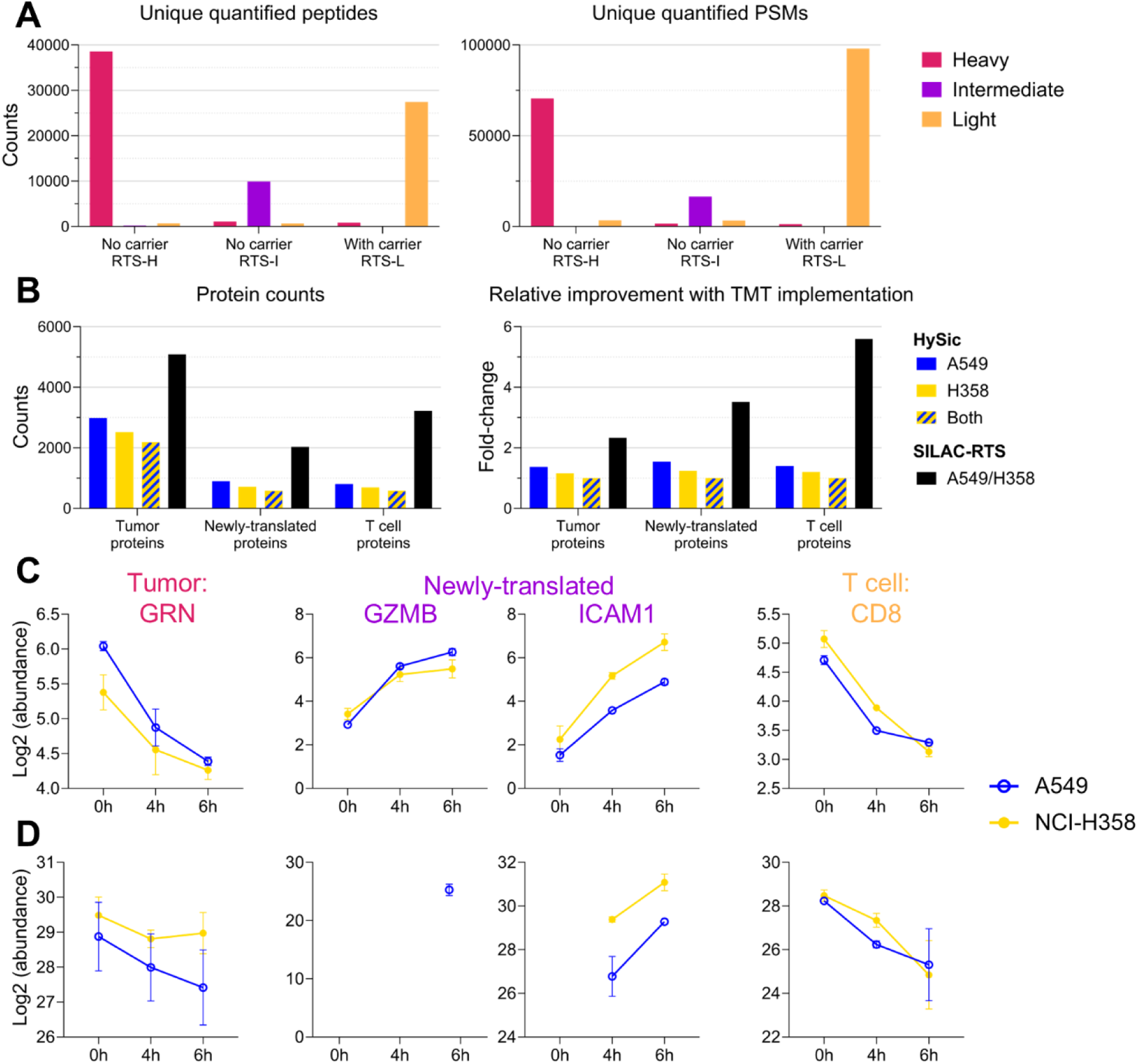
SILAC-RTS identifies more peptides and proteins than HySic, with a better temporal resolution. **(A)** Peptide (left) and PSM (right) counts in HySic samples prior to filtering through the analysis pipeline (Fig. S3B). **(B)** Protein counts in SILAC-RTS compared to the ones of HySic (left) and relative fold-change improvement (right). Proteins were counted in SILAC-RTS if they were measured with at least 2 PSMs. In HySic proteins found in at least 2 biological replicates in all 3 timepoints were considered. Measured intensities of key proteins with **(C)** SILAC-RTS and **(D)** HySic.

At the protein level, 5,086 heavy tumor proteins were quantified, 2,030 intermediate newly-translated proteins and 3,223 light T cell proteins (Fig 3B, left panel, black bars). Compared to the number of proteins found in both cell lines, in at least 2/3 biological replicates and all 3 timepoints in the original HySic analysis, this represents a 2.3-fold improvement for tumor protein coverage, 3.5-fold for newly-translated proteins and 5.6-fold for T-cell protein detection (Fig. 3B, right panel). Importantly, this impressive improvement in proteome coverage was achieved using less overall MS acquisition time.

To further validate our SILAC-RTS method, we examined key proteomics changes previously observed in the HySic study (10). Namely, for tumor proteins, progranulin (GRN) was confirmed as one of the proteins showing the most decreased abundance over time. Both granzyme B (GZMB) and intercellular adhesion molecule 1 (ICAM1) were validated as newly-translated and CD8A was found to be the most downregulated T-cell protein. TMT-based quantification not only reproduced these trends (Fig. 3C) but also provided superior resolution and reduced replicate variability(Fig. 3D). The near-absence of missing values in TMT data contributed to more accurate temporal profiling, especially for newly-translated proteins.

Our findings emphasize the importance and applicability of multiplexing SILAC-labeled co-culture samples using TMT to enhance proteome coverage, quantification accuracy and temporal resolution. This increased sensitivity may facilitate more detailed comparison across cell lines and experimental conditions.

### Differential proteomics signatures uncover divergent T cell response to A549 and NCI-H358

To test the hypothesis that the enhanced sensitivity could enable deeper insights into cell-cell interaction, we explored temporal changes in protein abundance across tumor cells and T cells during co-culture.

We began by examining significantly downregulated tumor proteins over time using hierarchical clustering (Fig. 4A). Despite being the category in which we identified the highest number of proteins, only a small number of significantly downregulated proteins were found (n = 42), suggesting limited co-culture-induced protein turnover in tumor cells within the first few hours of engagement. Among these, one cluster of proteins consistently more abundant in the T-cell resistant A549 than the sensitive NCI-H358 (cluster 1) was enriched in a pathway of disease of glycosylation (Fig. 4B).

**Figure 4.**
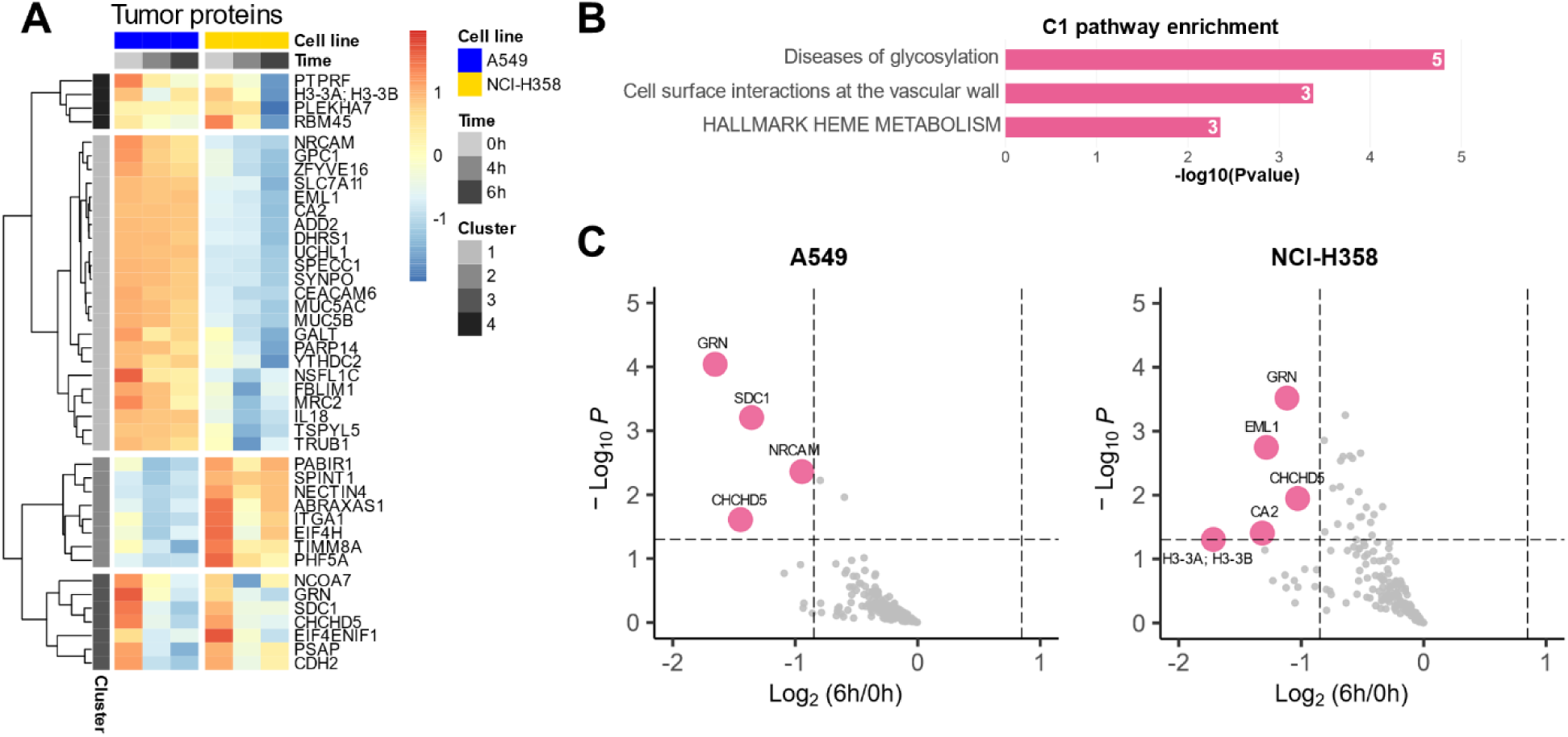
Protein changes in tumor. **(A)** Heatmap of significantly downregulated tumor proteins (n=42) in both A549 and NCI-H358 over time. Proteins with more than 1 missing value were discarded to allow heatmap clustering. Intensities are row scaled. **(B)** Metascape pathway enrichment in cluster 1 from panel (A). **(C)** Volcano plots comparing proteome profiles at 6 h versus 0 h in A549 (left) and NCI-H358 (right). Colored dots are significantly downregulated proteins at 6 h (adjusted p-value < 0.05) with a log_2_ fold change < −0.84.

When analyzing each cell line individually (Fig. 4C), both showed downregulation of progranulin (GRN) over time, consistent with previous findings (10). Notably, A549 also displayed reduced levels of cancer-associated proteins such as SDC1 and NRCAM.

We next focused on T cells, where 2,482 proteins were significantly downregulated during co-culture (Fig. 5A), indicating extensive protein turnover. Hierarchical clustering followed by pathway enrichment analysis revealed several biologically relevant terms (Fig. S5A). For example, cluster 3 which is characterized by higher baseline expression in NCI-H358 co-cultures, was enriched for JAK-STAT signaling, a pathway involved in immune response and interferon (IFN) γ production (28). However, the large number of downregulated proteins made it difficult to draw clear conclusions about T cell responses to each tumor cell line.

**Figure 5.**
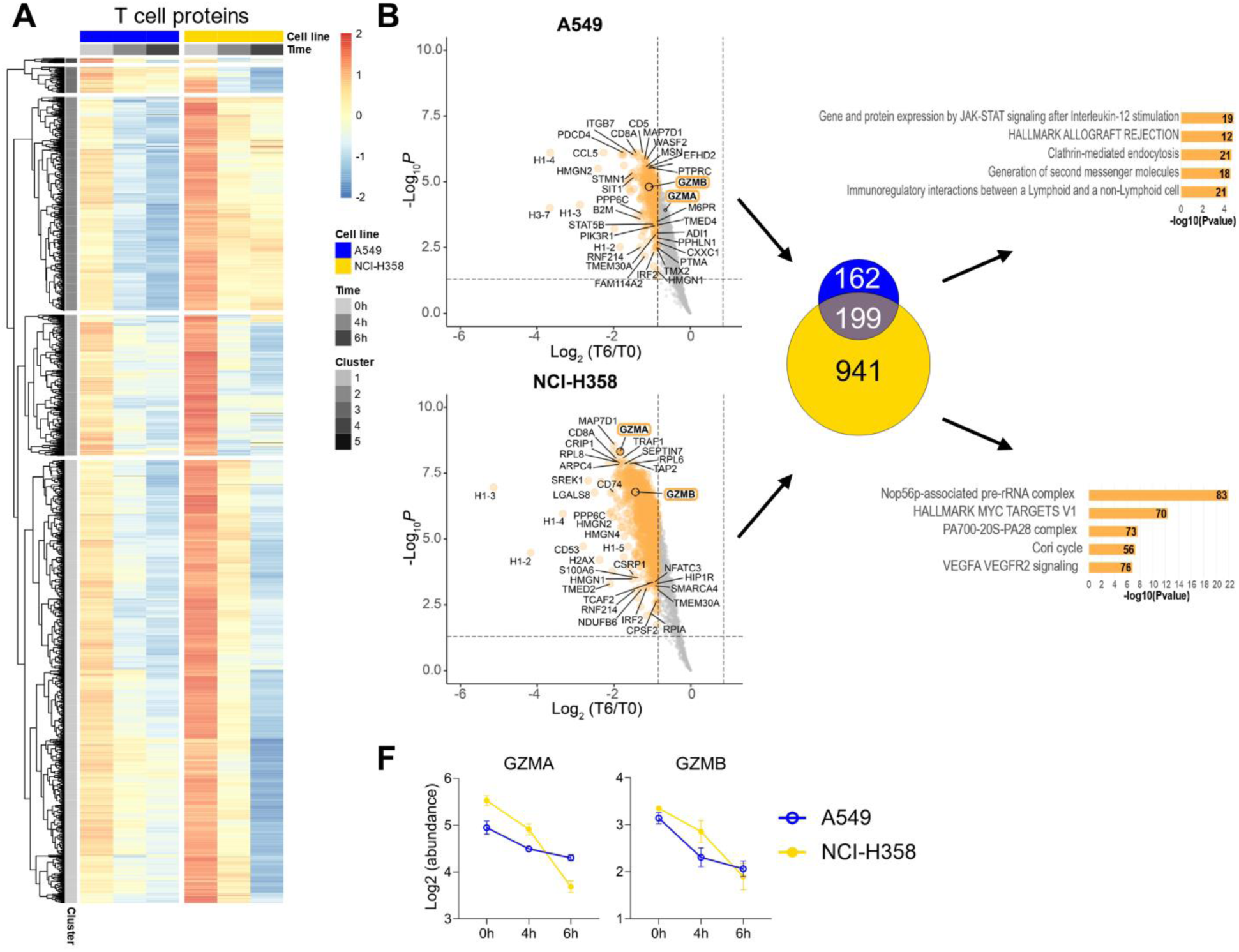
Protein changes in T cells. **(A)** Heatmap of significantly downregulated T cell proteins (n=2,482) in both A549 and NCI-H358 over time. Proteins with more than 1 missing value were discarded to allow heatmap clustering. Intensities are row scaled. **(B)** Volcano plots comparing proteome profiles at 6 h versus 0 h in A549 (top) and NCI-H358 (bottom). Proteins significantly downregulated at 6 h (adjusted p-value < 0.05 and log₂ fold change < −0.84) are highlighted and compared across cell lines using a Venn diagram. Separate metascape pathway enrichment analysis were performed for significantly downregulated proteins in A549 (top bar chart) and NCI-H358 (bottom bar chart). **(C)** Granzyme A (GZMA) and B (GZMB) intensities in both cell lines over time.

To address this, we investigated proteins significantly downregulated over time in T cells co-cultured with each tumor line separately (Fig. 5B, Fig. S5B, Fig. S5C). We then identified proteins uniquely downregulated in either A549 or NCI-H358 conditions and performed pathway enrichment analysis on these subsets, as well as on the shared proteins (Fig. S5D). In T cell co-culture with A549, uniquely enriched pathways were predominantly immune-related. These included “JAK-STAT signaling following interleukin-12 stimulation”, “second messenger generation” (critical for TCR signaling (29)), and “immunoregulatory interactions between lymphoid and non-lymphoid cells”, closely mirroring our experimental setup.

We observed nearly 6 times more unique downregulated proteins in T cells exposed to NCI-H358 as compared to A549. Pathways uniquely enriched in NCI-H358 co-cultures reflected a highly active T cell state. These consisted of the “Nop56p-associated pre-rRNA complex” (indicative of elevated ribosomal activity (30)), the “PA700–20S–PA28 complex” (a key element in proteasomal degradation (31)), and “MYC target genes” (associated with T cell metabolic reprogramming upon activation (32, 33)). Additional enrichment of the “Cori cycle” pathway, which involves lactate metabolism, further supports the presence of metabolically active, proliferative T cells known to produce lactate as a byproduct of increased glycolysis (34, 35).

Finally, we observed that granzyme B (GZMB) was among the most significantly downregulated proteins in T cells from both co-cultures, while granzyme A (GZMA) was only significantly downregulated in NCI-H358 (Fig. 5B, C), suggesting differential cytotoxic activity depending on the tumor context.

Together, these findings reveal distinct proteomic responses in tumor cells as well as T cells, depending on the tumor cell line they interact with. While T cells from A549 co-cultures are associated with immune signaling, NCI-H358 co-cultures elicit a more proliferative T cell phenotype, which may be indicative of a higher activation state or an enhanced rate of turnover. These differences underscore the importance of tumor context in shaping immune cell behavior and highlight the utility of the SILAC-RTS approach in dissecting cell-type–specific proteome dynamics.

### Newly-translated proteins reveal immune signatures and resistance mechanisms

To further test the sensitivity and resolution of our system, we focused on the least abundant protein population: the newly translated proteins. Unlike T cell proteins, these cannot be boosted with a carrier channel, making them a stringent test case for the sensitivity of our workflow.

First, we identified 96 significantly upregulated proteins over time and performed hierarchical clustering on them (Fig. 6A). Notably, ICAM1 formed its own distinct cluster and emerged as the most abundant newly-translated protein in both co-culture systems, consistent with our previous observations (10). However, its upregulation was more pronounced in the co-culture of T cells with NCI-H358 rather than with A549 (Fig. S6A). This observation aligns with the higher baseline expression of ICAM1 in NCI-H358 tumor cells compared to A549.

**Figure 6.**
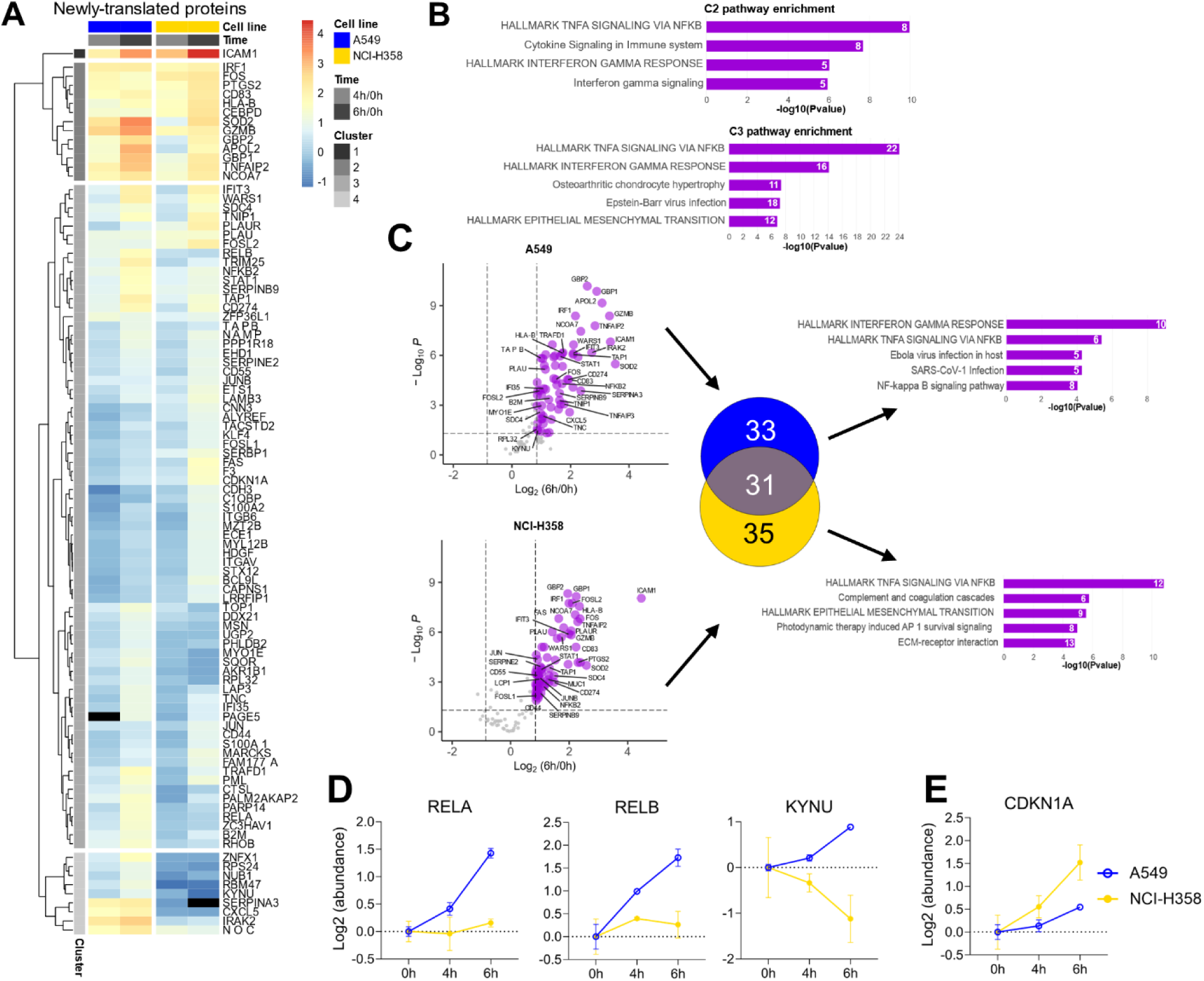
Changes in newly-translated proteins. **(A)** Heatmap of significantly upregulated newly-translated proteins (n=96) in both A549 and NCI-H358 over time. Proteins with more than 1 missing value were discarded to allow heatmap clustering. **(B)** Metascape pathway enrichment clusters 2 and 3 from panel (A). **(C)** Volcano plots comparing proteome profiles at 6 h versus 0 h in A549 (top) and NCI-H358 (bottom). Proteins significantly upregulated at 6 h (adjusted p-value < 0.05 and log₂ fold change > 0.84) are highlighted and compared across cell lines using a Venn diagram. Separate Metascape pathway enrichment analysis were performed for significantly upregulated proteins in A549 (top bar chart) and NCI-H358 (bottom bar chart). **(D-E)** Normalized intensities (relative to 0 h) of proteins contributing to the “Hallmark TNFA signaling via NFKB” pathway in panel (C) in A549 **(D)** and NCI-H358 **(E)**. Only proteins with established roles in cancer cell survival (D) or in cell cycle arrest and apoptosis (E) are shown.

Pathway enrichment analysis of the remaining clusters revealed strong immune-related signatures, particularly “Hallmark TNFA signaling via NFKB,” which was enriched in both clusters 2 and 3 (Fig. 6B). Interestingly, only cluster 3 also included a tumor-associated pathway: “Hallmark epithelial-mesenchymal transition.”

We next analyzed significantly upregulated newly-translated proteins over time in each co-culture system separately (Fig. 6C, S6B, S6C). A Venn diagram comparison revealed distinct and shared protein sets, which were subjected to pathway enrichment analysis (Fig. 6C). Again, both A549 and NCI-H358 co-culture conditions showed strong enrichment for “Hallmark TNFA signaling via NFKB.” We thus investigated the specific proteins driving this enrichment: 6 in A549 and 12 in NCI-H358. In A549, 3 out of the 6 proteins, namely RELA, RELB and KYNU, have documented roles in promoting cancer cell survival (Fig. 6D) (36–39). In contrast, 1 out of the 12 proteins in NCI-H358, CDKN1A, is known to mediate cell cycle arrest and apoptosis (Fig. 6E) (40). The remaining proteins in both conditions have dual roles in tumor biology (Fig. S6E, F). We also examined proteins commonly upregulated in both co-cultures. These were again enriched for “Hallmark TNFA signaling via NFKB” and other immune-related pathways (Fig. S6D), further reinforcing the predominance of immune signaling in the newly-translated proteome.

Finally, we observed that several proteins relating to immune evasion, CD274 (PD-L1), JAK1 (which stabilizes PD-L1 expression (41)) and SERPINB9 (which protects against granzyme B (42)) were more strongly upregulated in the co-culture with A549. This could suggest a more robust immune resistance phenotype in A549, potentially driven by the observed elevated levels of RELB and KYNU, both of which have previously been linked to heightened PD-L1 expression and immune evasion in other cancer models (43, 44). Overall, the analysis of newly-translated proteins highlights distinct immune-related responses and possibly different resistance mechanisms. While both A549 and NCI-H358 co-culture systems activate TNFA–NFKB signaling, A549 is more strongly associated with pro-survival and immune evasion pathways, including PD-L1 stabilization and granzyme B inhibition. These findings underscore the power of the SILAC-RTS approach to resolve subtle but biologically meaningful differences in low-abundance proteome dynamics.

## Discussion

In this study, we introduce a novel application of RTS, based on SILAC labeling to efficiently deconvolute individual proteomes within co-culture samples. Each cell type (tumor cells, T cells) and newly-translated proteins harbored distinct SILAC labels and co-culture samples were multiplexed with TMT. We also included a carrier channel in our TMT sample design to boost identifications of T cell proteins, which are less abundant than those of tumor cells.

SILAC-TMT combinations have previously been used in pulsed SILAC experiments to study protein translation (4) as well as in time-resolved analysis of proteome dynamics (14, 16). But to our knowledge, the integration of SILAC-based RTS has only been implemented once (45), and in that case, SILAC was used for enhanced sample multiplexing. The approach used here introduces an innovative and robust framework for the investigation of intercellular communication dynamics at the proteome level.

Beyond protein identification, we evaluated the quantitative accuracy of our method using a mixed sample containing equal amounts of two SILAC-labeled populations. Expectedly, our RTS-SILAC approach identified nearly as many proteins as MS^2^-based methods for each population and achieved MS^3^-grade quantitative accuracy (20, 26, 46). However, even in this ideal 1:1 peptide mix, the measured ratios across TMT channels did not fully match those obtained from each population individually. This discrepancy was unexpected, as standard mixed-species samples of yeast/human typically yield accurate quantification. We can speculate that the increased complexity of mixing two human proteomes (3x more diverse than the one of yeast) may increase the risk of co-isolation of peptides at the MS^1^ level, therefore lowering quantification accuracy.

Co-culture samples from the HySic study were originally performed with a cell ratio of 1:1 immune to cancer cells. Due to their smaller size T cells contain less protein (12, 13), leading to a protein ratio of 1:5 or 1:6 immune-to-cancer protein amounts. To characterize our method in conditions that mimic co-culture, we designed a mixed sample with two SILAC-labeled populations with a 1:5.5 ratio. This revealed that quantitative accuracy reduces for the less abundant population, highlighting the importance of representative sample design when evaluating method performance. Although the addition of a carrier channel increased protein identifications by 2.5 to 3.6-fold, it also worsened ratio compression, a known issue particularly pronounced at the MS^2^ level (17, 47). Nevertheless, with stringent data filtering, we were able to detect proteome changes specific to each SILAC label.

Strikingly, the T cell proteome exhibited extensive turnover, with nearly 80% of identified proteins (2,482 out of 3,223) significantly downregulated over time. This is substantially higher than the previously reported 7–9% turnover in naïve T cells responding to antigens (48), but aligns with the short half-life of certain T cell proteins (49) and the fact that our T cells were pre-activated with anti-CD3 and anti-CD28 antibodies (1). In contrast, less than 1% of tumor proteins (42 out of 5,086) were significantly downregulated, reflecting the slower turnover (average ∼20h for HeLa cells) and longer half-lives typical of tumor proteins (up to several days in pancreatic ductal adenocarcinoma organoids) (50, 51).

Pathway enrichment analysis uncovered distinct T cell activation states depending on the tumor cell line they were interacting with. T cells co-cultured with A549, which is resistant to T cell killing, showed downregulation of immune-related pathways, including downstream TCR signaling and immune regulation, suggesting a slow and subdued activation. Conversely, T cells exposed to sensitive NCI-H358 cells were enriched for pathways associated with several aspects of robust T cell activation and MYC target genes, which are typically induced in later activation phases following TNFα signaling via NF-κB (32, 52). The presence of the Cori cycle pathway, indicative of lactate metabolism and potential T cell exhaustion (53), further supports this interpretation. However, the enrichment of these pathways in the most downregulated proteins could imply either active engagement of the pathways, leading to high protein turnover, or instead their inhibition through protein degradation. Additional studies would be required to determine which of these mechanisms is at play.

Consistent with the observed high turnover of T cell proteins, newly-translated proteins were enriched for immune-related pathways, particularly “Hallmark TNFA signaling via NFKB” in both co-culture systems. TNFα signaling is an early hallmark of T cell activation (52, 54) and can promote either survival or apoptosis. Interestingly, newly-translated proteins in the A549 co-culture included several pro-survival factors, while those in the NCI-H358 co-culture featured one protein involved in cell cycle arrest. TNFAIP3 (A20), a key TNF regulator implicated in survival vs death signaling, appeared higher in A549 (55). Of note, TNFAIP3 was not detected in NCI-H358 at 0 h, and therefore 0 h-normalized data are not shown in Fig. S6E. While experimental validation would be required to determine the cellular origin of these proteins, the high turnover in T cells suggests they may be T cell-derived. The significant upregulation of RELA and RELB in the A549 co-culture may indicate activation of pro-survival pathways in activated T cells (56, 57), whereas increased KYNU expression could reflect either enhanced T cell activity or exhaustion (58, 59).

Among the most downregulated proteins in A549 were SDC1 and NRCAM. Despite its downregulation, NRCAM remained more abundant in A549 than in NCI-H358 (not shown) and is known to promote metastasis (60). SDC1 inhibition, on the other hand, facilitates CD8 T cell infiltration into tumors (61). These findings, along with the enrichment of the “disease of glycosylation” pathway, driven by mucins MUC5AC and MUC5B, in the cluster characterized by higher expression in A549, suggest that A549 cells may be less accessible to T cells. Mucins form gel-like barriers that shield cells from immune attack (62), and MUC5AC has been linked to angiogenesis, metastasis, and immune evasion in lung adenocarcinoma (63). Alteration of glycosylation-related proteins may thus contribute to physical shielding and impaired immune synapse formation, enhancing tumor survival under immune pressure. Supporting this, ICAM1, a key adhesion molecule for T cell-tumor synapse formation, was more abundant and upregulated in NCI-H358, potentially explaining its greater sensitivity to T cell-mediated killing (64).

Further analysis revealed that T cells interacting with NCI-H358 employed both caspase-dependent (GZMB) and caspase-independent (GZMA) killing pathways, whereas those exposed to A549 relied solely on caspase-dependent mechanisms (65). Additionally, the immune checkpoint PD-L1 and its stabilizer JAK1 were more highly translated in the A549 co-culture (41), along with SERPINB9, a granzyme B inhibitor. Although the origin of these proteins is uncertain, literature suggests they likely derive from tumor cells and contribute to immune evasion (27).

The implementation of TMT into the HySic workflow not only dramatically enhanced proteome depth but it also offered the added benefit of requiring less MS time. For the 18 samples investigated in the present study, DDA MS^1^-based precursor quantification previously required approximately 52 h (175 min runs), whereas TMT multiplexing combined with high-pH fractionation allowed all RTS acquisitions (L, H, I) to be completed in just about 50 h.

In summary, our SILAC-RTS method enables detailed analysis of co-culture systems by leveraging RTS and SILAC to distinguish cell populations. We have also demonstrated the benefit of a carrier channel to enhance signal detection and to provide deeper insights into intercellular communication. Many of the findings presented here, while requiring further validation, were previously obscured by MS^1^-based precursor quantification. SILAC-RTS offers superior resolution of protein-level trends and serves as a powerful hypothesis-generating tool for studying communication dynamics at the proteome level.

## Abbreviations

ACN: acetonitrile
DDA: data-dependent acquisition
FDR: false-discovery rate
TMT: tandem mass tag
LC-MS/MS: liquid chromatography coupled to tandem mass spectrometry
MS: mass spectrometry
NSCLC: non-small cell lung cancer
PSM: peptide-spectrum match
RTS: real-time search
RTS-H: real-time search with heavy SILAC as a fixed modification
RTS-I: real-time search with intermediate SILAC as a fixed modification
RTS-L: real-time search with light SILAC as a fixed modification
SILAC: stable isotope labeling by amino acids in cell culture
S/N: signal-to-noise
SPS: synchronous precursor selection
TMT: tandem mass tags

## Acknowledgment

We would like to thank Esther Nolte-’t Hoen and Marije Kuipers for providing the heavy-labeled PC3 samples.

## Author contribution

Conceptualization: A.C., K.S., M.A.; Methodology: A.C., K.S.; Validation: A.C.; Formal analysis: A.C.; Investigation: A.C.; Resources: S. I-M, K.S., D.S.P; Data curation: A.C.; Writing – Original Draft: A.C.; Writing – Review & Editing: A.C., K.S., M.A.; Visualization: A.C.; Supervision: K.S.; Funding acquisition: M.A., D. S. P

## Data availability

All mass spectrometry proteomics data have been deposited to the ProteomeXchange Consortium via PRIDE with the dataset identified PXD067452.

## Supplemental data

This article contains supplemental data (figures and tables).

