## Supplementary material for "Decoding interaction-induced proteome changes in co-cultures with hybrid quantification and SILAC-directed real-time search": SI figures

**Table of contents**

- Supplemental Fig. S1
- Supplemental Fig. S2
- Supplemental Fig. S3
- Supplemental Fig. S4
- Supplemental Fig. S5
- Supplemental Fig. S6

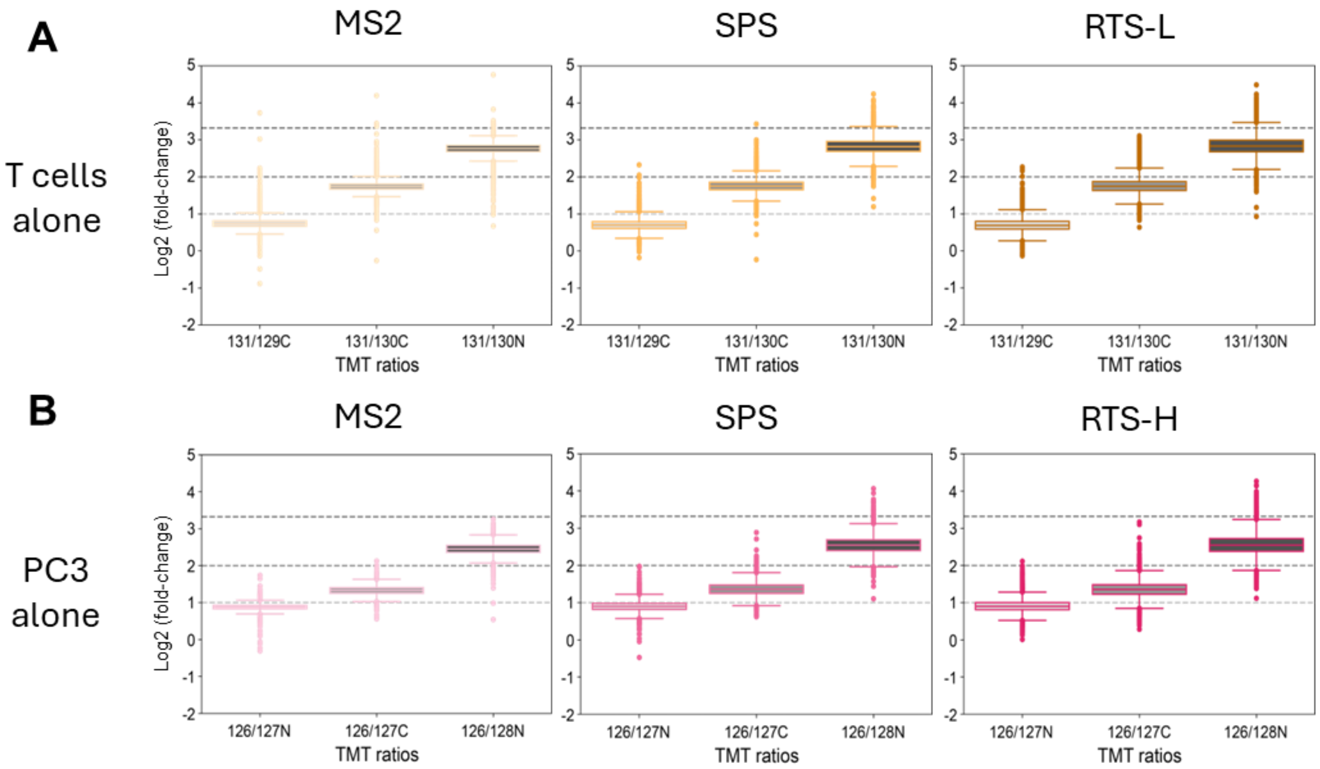


**Figure S1. Peptide-level quantification.** Distribution of quantified **(A)** light and **(B)** heavy spectra using TMT ratios across relevant channels. Expected theoretical ratios are marked by horizontal dashed lines.

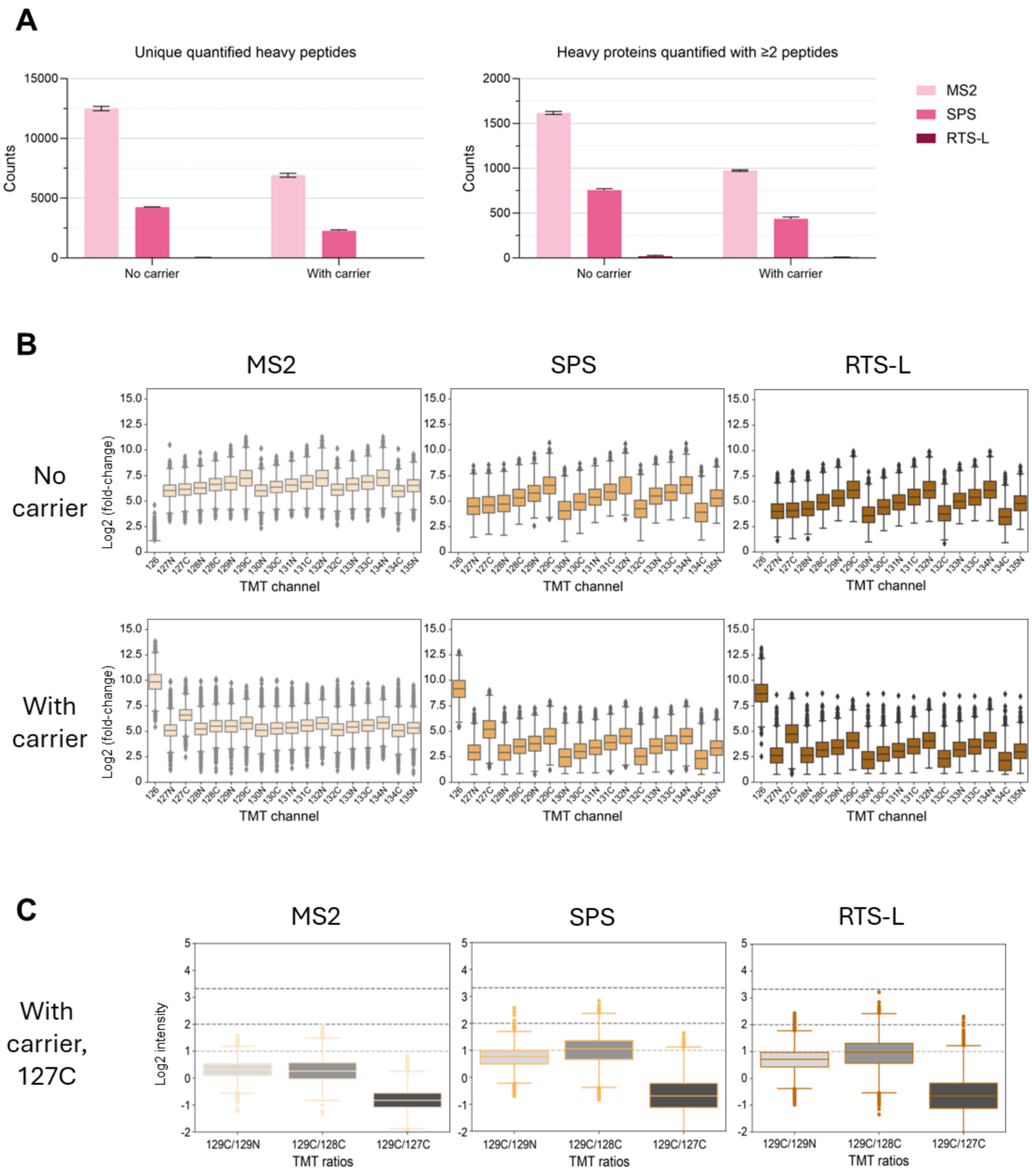


**Figure S2. Effect of the carrier channel on heavy peptide counts, peptide intensities and accuracy. (A)** Heavy peptide and protein counts across MS acquisitions in the absence or presence of a carrier channel. **(B)** Distribution of light peptide intensities across TMT channels in all MS acquisitions, in the presence (top) or absence (bottom) of a carrier channel. **(C)** Distribution of quantified light peptides in the presence of a carrier channel. TMT ratios across relevant channels and across MS acquisitions are represented. Expected ratios are shown by the horizontal dashed lines.

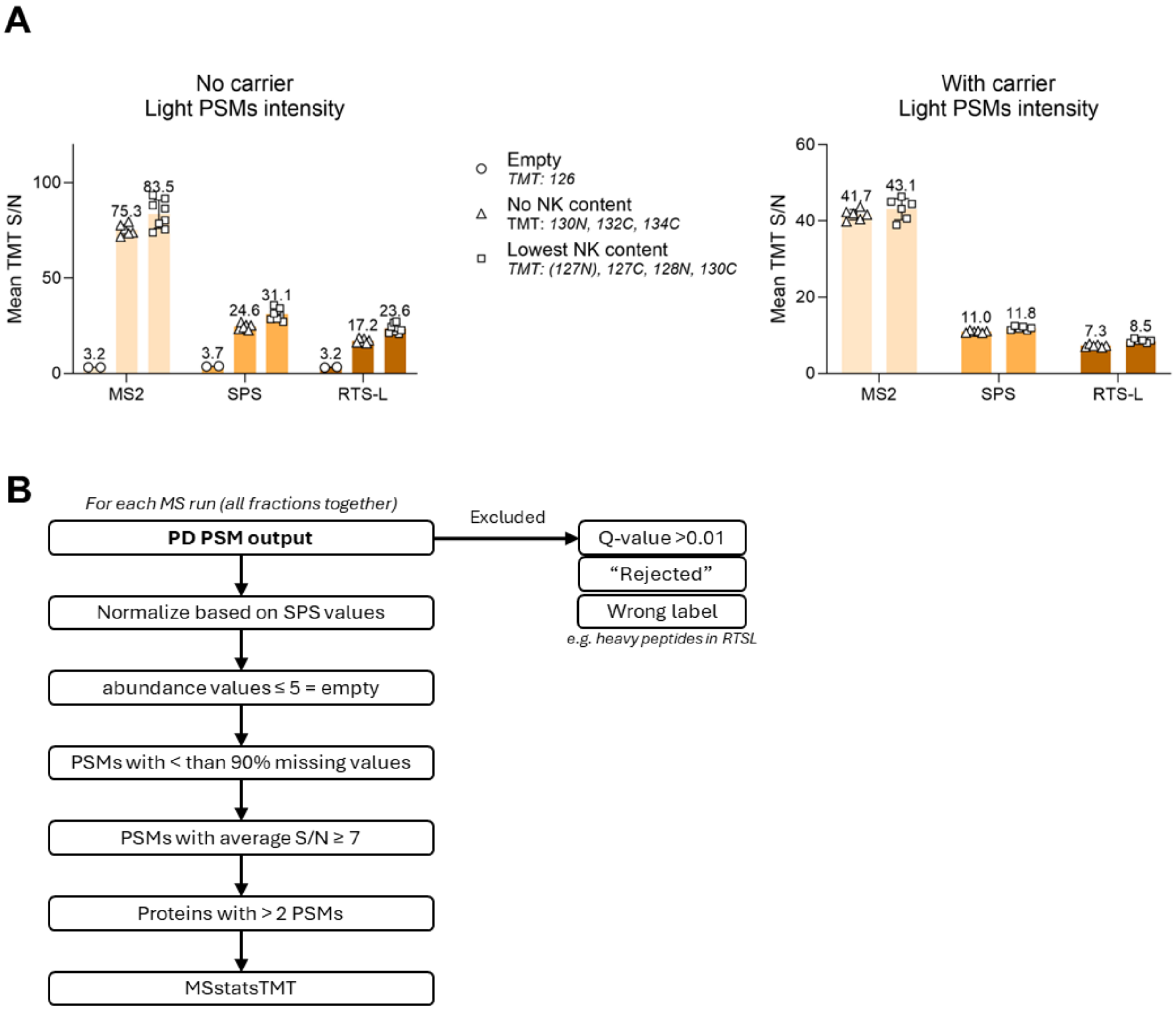


**Figure S3. TMT noise level in mixed SILAC samples. (A)** Mean TMT intensities at the PSM-level in the absence (left) or presence (right) of a carrier channel in channels containing no peptides, no light NK peptides or containing the lowest amount of NK light peptides. **(B)** Analysis pipeline implementing TMT noise cutoffs.

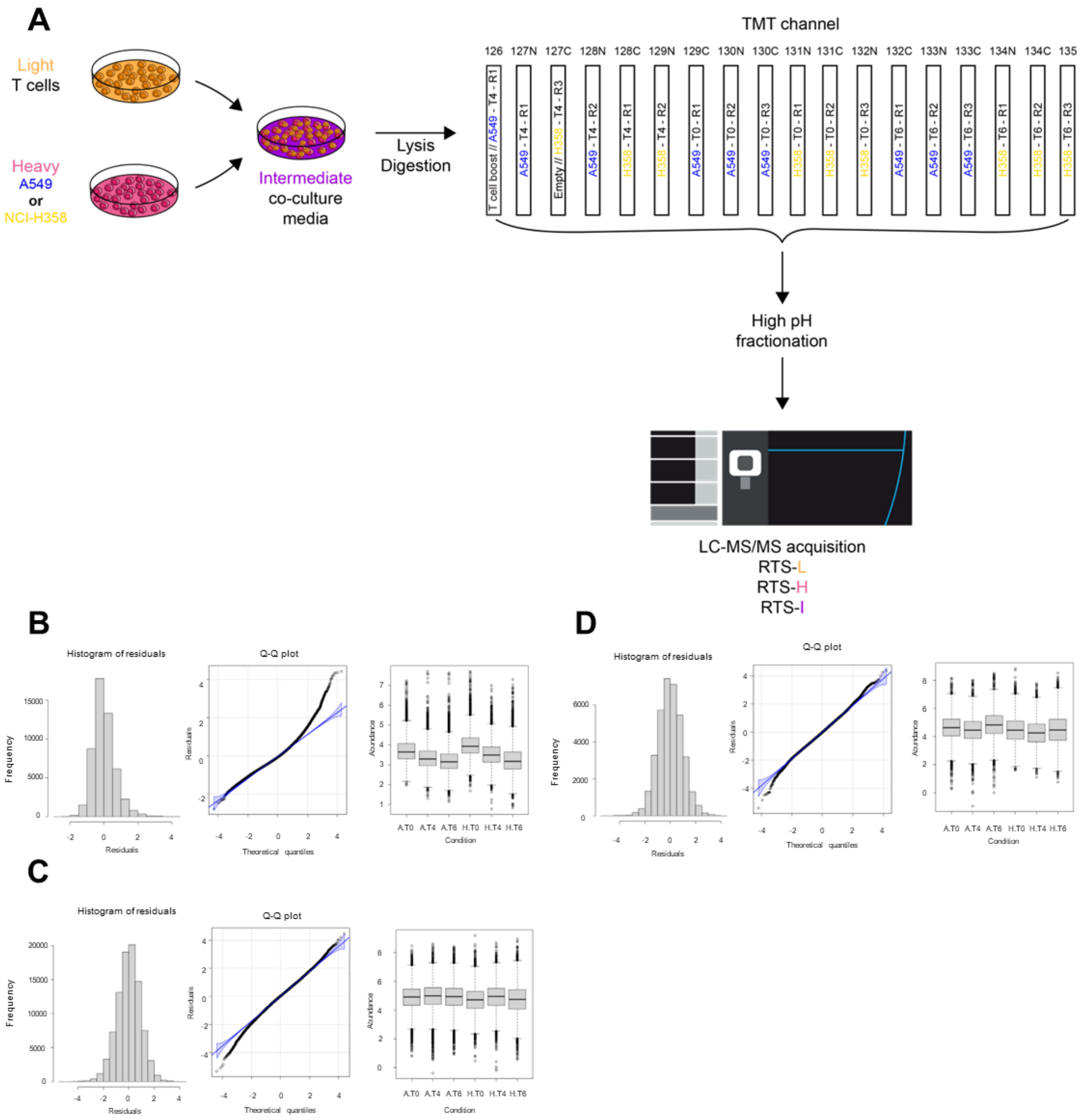


**Figure S4. HySic-SILAC-RTS sample design and normality. (A)** Previously, light primary T cells were co-cultured with heavy-labeled A549 or NCI-H358 tumor cells in a medium containing intermediately-labeled amino acids. Co-culture samples were collected altogether, lysed and digested. Digests were then TMT labeled, fractionated with high-pH LC and measured on an Orbitrap Eclipse with 3 separate acquisition methods using each SILAC labels as a trigger for the acquisition of $MS^3$ quantitative scans. **(B-D)** Assessment of normality of light **(B)**, heavy **(C)** and intermediate **(D)** datasets with distribution (left) and QQ plot (middle) on the residuals and abundance distribution (right).

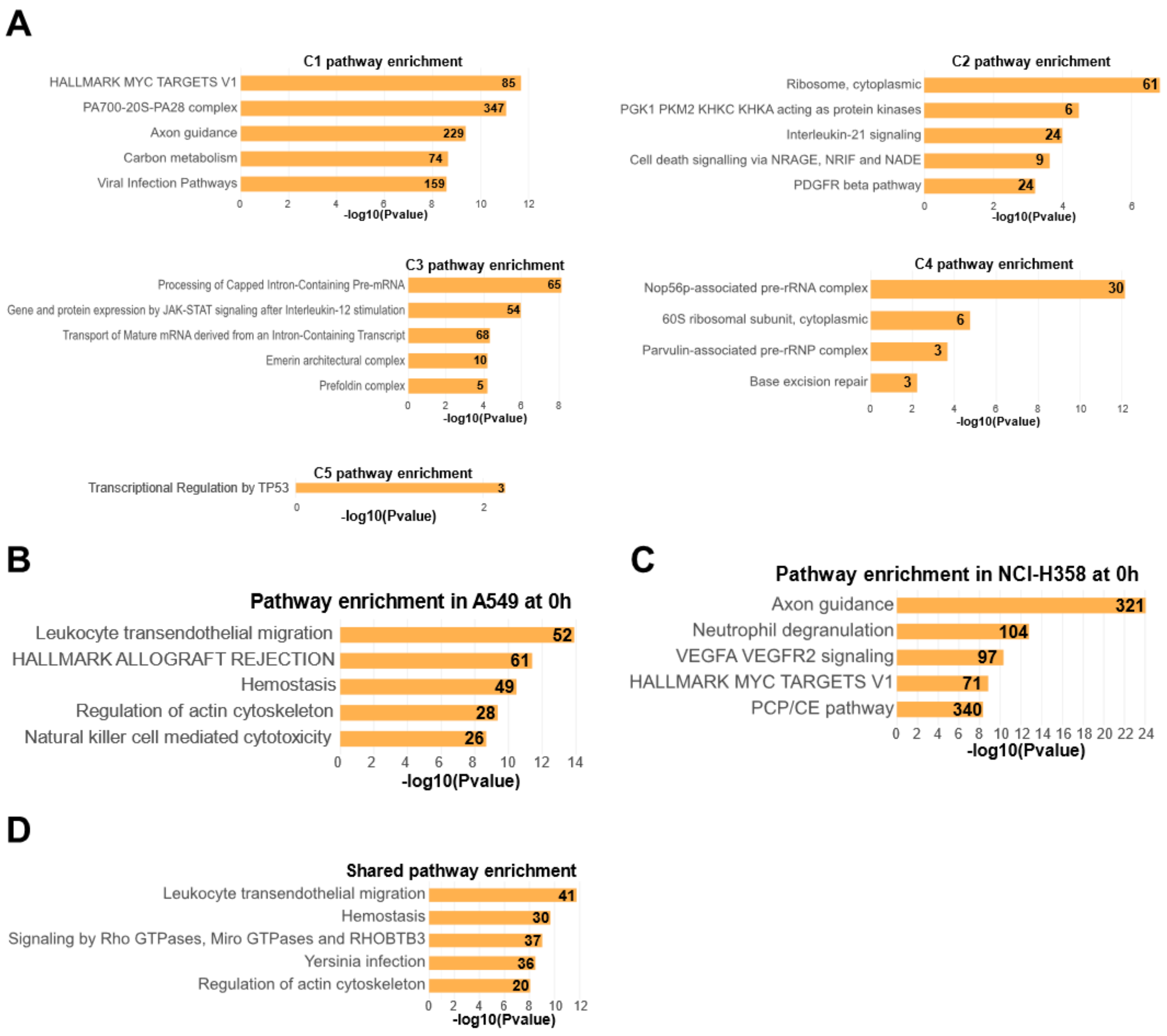


**Figure S5. Pathway enrichment analyzes in tumor and T cells. (A)** Metascape pathway enrichment in clusters 1, 2, 3, 4 and 5 from Fig. 4D. **(B-C)** Metascape pathway enrichment on significantly downregulated proteins identified in Fig. 4E in A549 **(B)** and in NCI-H358 **(C)**. **(D)** Metascape pathway enrichment in significantly downregulated proteins found in both cell lines (intersection of Venn diagram in Fig. 4E).

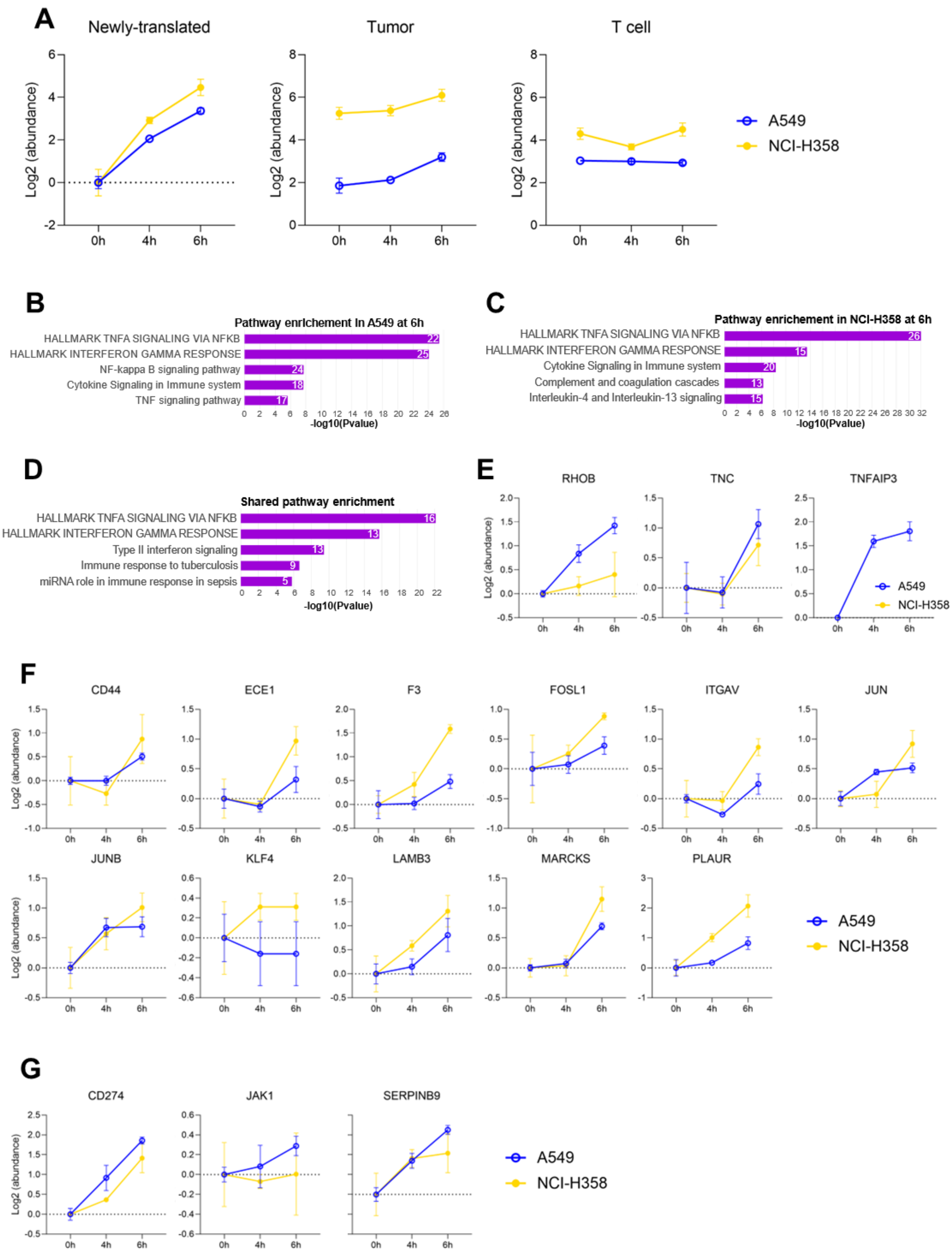
A
Newly-translated
Tumor
T cell
Log2 (abundance)
0h
4h
6h
A549
NCI-H358
B
Pathway enrichment in A549 at 6h
HALLMARK TNFA SIGNALING VIA NFKB
HALLMARK INTERFERON GAMMA RESPONSE
NF-kappa B signaling pathway
Cytokine Signaling in Immune system
TNF signaling pathway
-log10(Pvalue)
C
Pathway enrichment in NCI-H358 at 6h
HALLMARK TNFA SIGNALING VIA NFKB
HALLMARK INTERFERON GAMMA RESPONSE
Cytokine Signaling in Immune system
Complement and coagulation cascades
Interleukin-4 and Interleukin-13 signaling
-log10(Pvalue)
D
Shared pathway enrichment
HALLMARK TNFA SIGNALING VIA NFKB
HALLMARK INTERFERON GAMMA RESPONSE
Type II interferon signaling
Immune response to tuberculosis
miRNA role in immune response in sepsis
-log10(Pvalue)
E
RHOB
TNC
TNFAIP3
F
CD44
ECE1
F3
FOSL1
ITGAV
JUN
JUNB
KLF4
LAMB3
MARCKS
PLAUR
G
CD274
JAK1
SERPINB9

**Figure S6. Characterization of newly-translated proteins. (A)** ICAM1 protein intensity in newly-translated (left), tumor (middle) and T cells (right) in both cell lines. **(B-C)** Metascape pathway enrichment on significantly upregulated proteins identified in Fig. 5C in A549 **(B)** and in NCI-H358 **(C)**. **(D)** Metascape pathway enrichment in significantly upregulated proteins found in both cell lines (intersection of Venn diagram in Fig. 5C). **(E-F)** Normalized intensities (relative to 0 h) of proteins contributing to the "Hallmark TNFA signaling via NFKB" pathway in Fig. 5C in A549 **(E)** and NCI-H358 **(F)**. **(G)** Normalized intensities (relative to 0 h) of proteins with an established role in tumor resistance to immune killing.
